# The molecular prognostic score, a classifier for risk stratification of high-grade serous ovarian cancer

**DOI:** 10.1101/2023.06.19.545525

**Authors:** Siddik Sarkar, Sarbar Ali Saha, Poulomi Sarkar, Sarthak Banerjee, Pralay Mitra

**Affiliations:** Cancer Biology & Inflammatory Disorder, Translational Research Unit of Excellence (TRUE), CSIR-Indian Institute of Chemical Biology, Kolkata, 700032, WB, INDIA; Academy of Scientific and Innovative Research (AcSIR), INDIA; Computer Science and Engineering, Indian Institute of Technology Kharagpur, Kharagpur, 721302, WB, INDIA

**Author notes:** Corresponding Author: Siddik Sarkar, CSIR-Indian Institute of Chemical Biology.

**Keywords:** High grade serous ovarian cancer (HGSOC), Prognosis, molecular prognostic score (mPS), Biomarker, Over-all survival (OS)

## Abstract

The clinicopathological parameters such as residual tumor, grade, FIGO score are often used to predict the survival of ovarian cancer patients, but the 5-year survival of high grade serous ovarian cancer (HGSOC) still remains around 30%. In recent years, a gene expression based molecular prognostic score (mPS) was developed that showed improved prognosis in several cancers including ovarian cancer.

The feature extraction using LASSO-Cox regression was applied on the training data with 10-fold cross validation to obtain 20 predictor genes along with the coefficients to derive mPS. The mPS based prognosis of HGSOC patients was validated using the log-rank test and receiver operator characteristic curve.

The AUC of 20 gene-based mPS in predicting the 5-year overall survival was around 0.7 in both the training (n=491) and test datasets (n=491). It was also validated across HGSOC patients (n=7542), data collected from the Ovarian Tumor Tissue Analysis (OTTA) consortium. The mPS showed significant impact (adjusted HR = 6.1, 95% CI of HR= 3.65-10.3; p <0.001) on prognosis of HGSOC. The performance of mPS for the prognosis of survival of HGSOC was substantially better than conventional parameters: FIGO (adjusted HR=1.1, 95% CI=0.97-1.2, p=0.121), residual disease (adjusted HR=1.3, 95% CI= 1.13-1.4, p<0.001), and age (adjusted HR=1.2, 95% CI= 0.98-1.6, p=0.08). It was found that focal-adhesion, Wnt and Notch signaling pathways were significantly (p<0.001) upregulated, whereas antigen processing and presentation (p<0.001) was downregulated in high risk HGSOC cohorts based on mPS stratification.

The molecular prognostic score derived from 20-gene signature is found to be the novel robust prognostic marker of HGSOC. It could potentially be harnessed in clinical settings to determine the overall survival of ovarian cancer. The high risk HGSOC patients based on mPS stratification could be benefited from alternative therapies targeting Wnt/ Notch signaling pathways and also immune evasion.

**Author summary:** The 20-gene signature based molecular prognostic score (mPS) was found to be associated with risk stratification and hence, predicting the overall survival time of HGSOCs. It was applicable in all training HGSOC datasets and across RNA sequencing platforms that also includes the previous reported studies in HGSOC cohorts (Millstein et al., 2020; Talhouk et al., 2020). The 20-gene signature based mPS for the prognosis of overall survival of HGSOC outperformed conventional parameters: age, residual disease and FIGO score. The high or increased risk group of HGSOC based on our mPS stratification was found to have dysregulated pathways of Wnt, Notch, Akt signaling, and antigen presentation. Thus, treatments targeting these pathways might be beneficial for high risk HGSOC and hence anticipated to improve the over-all survival of HGSOC.

## Introduction

Epithelial ovarian cancer (EOC) is classified into different categories based on histotypes and grade [1]. Despite the initial responses with cytoreductive surgery and platinum based chemotherapy, high grade serous ovarian cancer (HGSOC) continued to account for 70% of EOC-related cases with more than 75% death within 10 years of initial diagnosis. It might be due to the high rate of intra-tumor genetic heterogeneity and chromosome instability within the HGSOCs [2, 3] subsequently supporting clonal evolution [4], resulting in chemo- or therapy-resistance. Therefore, a search for efficient gene signatures or prognostic markers is an urgent unmet clinical need for HGSOC.

Survival prediction takes various factors into account; like age, FIGO stage, histology, residual disease and tumor recurrence [5, 6]. However, prediction based on these orthodox clinical information has limited potential to give rise to a robust prognostic method. It is because of the complex interaction of various molecules as well as immunological factors leading to variable responses within the HGSOCs. Recently the molecular subtypes of HGSOCs based on transcriptome profiles have been identified [7, 8]. The most common and consensus subtypes using various clustering algorithms are mesenchymal, immunoreactive, differentiated and proliferative. Although these molecular subtypes showed distinct and differential regulation of biological pathways between the groups, but showed relatively less influence on the survival of patients using TCGA HGSOC cohort data [9]. It has been reported previously that gene signatures could potentially and significantly played role in determining the survival of cancer patients [10] including ovarian cancer [9]. A similar approach has been applied using 101-prognostic gene signatures for predicting the survival of HGSOCs [11]. This approach of using molecular gene signatures as prognostic marker has been studied or reported in various cancers: breast [12], colon [13] and prostate [14].

Herein, we proposed to develop a molecular prognostic score (mPS), a machine learning approach for stratifying the prognosis of HGSOCs based on the expression of only 20 predictor genes and the associated coefficients as derived from LASSO-Cox regression [15]. The proposed study design was schematically shown in Fig. 1. In this study, we have considered 1022 subjects/samples and screened or considered only 10225 genes that are found common in the Cancer Genome Atlas (TCGA) and Gene Expression Omnibus (GEO) databases. The micro-array based expression analysis of the same or similar platform (Affymetrix human U133A microarray or Affymetrix Human Genome U133 plus 2.0 Array) has been used here to filter the common genes for subsequent analysis. These common genes across different datasets were further screened to obtain prognostic gene signature of HGSOCs based on Cox (proportional hazards) regression model [16]. Finally, further trimming of prognostic genes and feature extraction was done by applying the LASSO Cox regression model [16] on training datasets of HGSOCs. This resulted in obtaining predictive markers along with derived coefficients that were subsequently used to obtain mPS that eventually determined the prognosis in test or validation datasets (Fig. 1).

**Figure 1:**
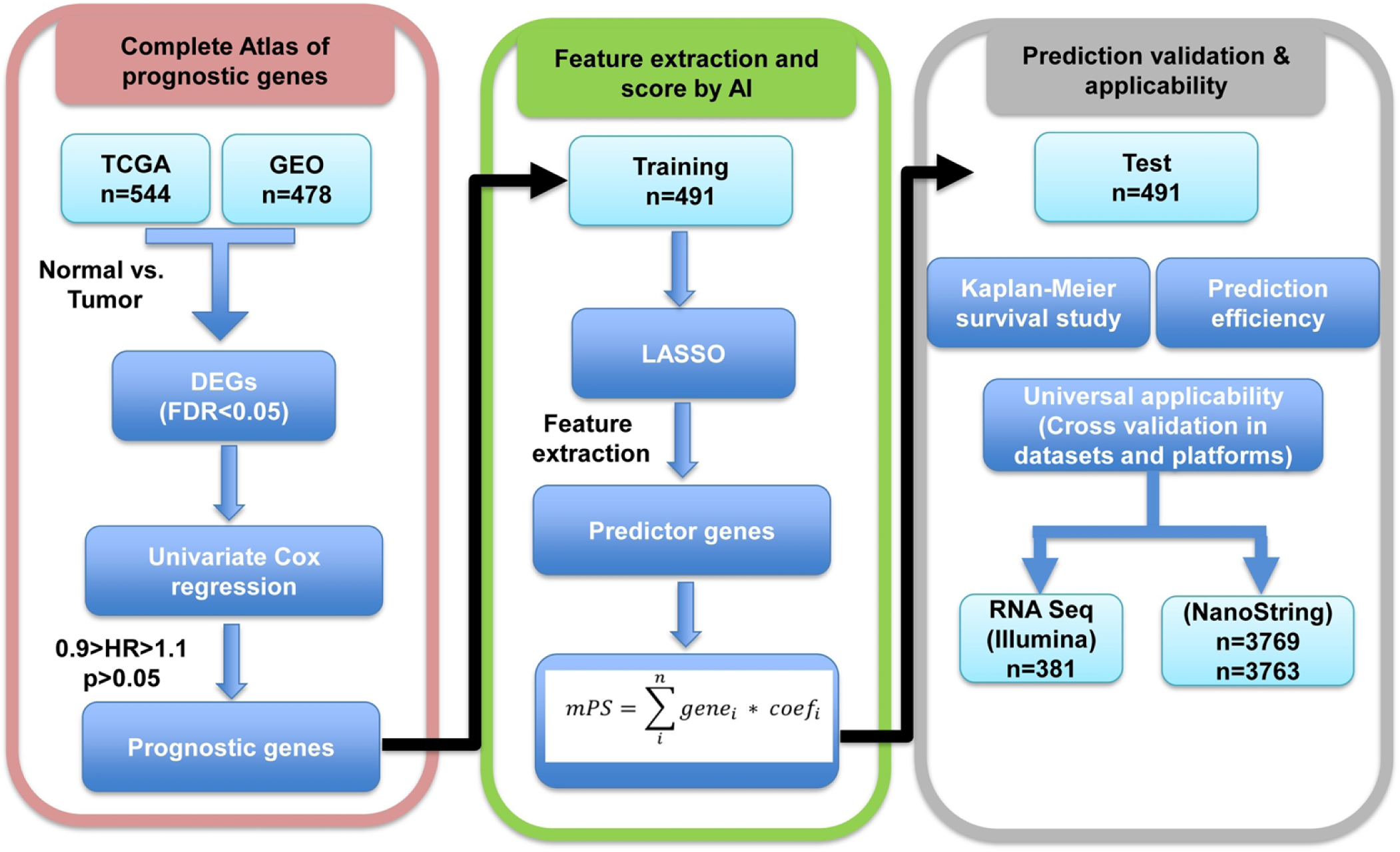
Methodology adapted to screen and filter genes for obtaining molecular Prognostic score (mPS) based on prognostic signature genes: RNA expression data obtained from TCGA and GEO as indicated were used to find prognostic genes. These prognostic genes were further used in training datasets, 10-fold cross validation to obtain predictor genes and associated coefficients (feature extraction) after applying LASSO regression. These predictor genes and the derived mPS were applied in validation or test datasets. It was also applied in different mRNA expression platforms such as RNA Sequencing by Illumina and NanoString. DEGs; Differentially expressed genes that are significantly (FDR *<*0.05) expressed between tumor samples as compared to normal samples. AI; Artificial intelligence. OTTA-SPOT; Ovarian Tumor Tissue Analysis consortium - Stratified Prognosis of Ovarian Tumors.

## Methods

### Datasets

Gene expression raw microarray dataset were downloaded from the cancer genome atlas (TCGA) viz., TCGA-OV (https://gdac.broadinstitute. org/?cohort=OV) and Gene Expression Omnibus (GEO) managed by the National Center for Biotechnology Information (NCBI)(https://www.ncbi. nlm.nih.gov/geo/). The GEO datasets are GSE18520 (n=63), GSE26712 (n=195), GSE26193 (n=79), GSE63885 (n=73), GSE14764 (n=68). Both the TCGA-OV (n=544) and GEO datasets (n=478) accounting 1022 as a total number of clinical samples and 10225 as the common gene symbols found in all datasets. The individual datasets were processed and normalized using the Robust Microarray Average (RMA) approach. Further quantile normalization (normalization between arrays) followed by removal of batch effect was performed between different datasets to have a similar pattern or log-ratios of similar distributions across various datasets (Appendix A Suppl Fig. S1). To rule out/ eliminate the outliers in the samples, a correlation matrix of mRNA expression of samples (Array-Array Intensity correlation) [17] was performed. We have used a correlation cut-off 0.7among the samples/ subjects. So, samples/ subjects (n=1016) with a correlation ≥ 0.7 is taken into consideration for subsequent analysis.

Differential gene expression was performed between HGSOC (n=988) cases vs. control sample (n=28) using R (version 4.1.0)/ Bioconductor, limma, and several associated packages. The fold-change (FC) ≥1.5 and false discovery rate (FDR) *<*0.05 was used for studying or selecting the differential gene expression. The detailed methodology was schematically shown in Fig. 1.

### Univariate analyses on differential gene expression

The significant (FDR*<*0.05) <u>d</u>ifferential <u>e</u>xpressed <u>g</u>enes (DEGs) between HGSOC tumor vs control as explained above were selected. To study prognostic genes, univariate cox proportional hazards regression analyses [16] was applied using these differential expressed genes (HGSOC vs. Control) and the associated survival data of HGSOC cohorts. The genes that played a role in the survival of HGSOC patients were further filtered by applying log-rank p-value *<*0.05 and 0.9*>*hazard ratio (HR) *>*1.1. This similar strategy of pre-filtering was applied previously[13, 18] prior to multivariate analyses to reduce noise (number of genes *>>* number of samples) [12, 19].

### Regularized Cox Regression on selected genes based on univariate cox genes

The selected genes obtained using univariate analyses were further used to conduct a multivariate regression analysis. Here we had applied a least absolute shrinkage and selection operator (LASSO) estimation using R/ Rstudio with package “glmnet” [15, 20]. The HGSOC samples were divided randomly into training (n=491) and test (n=491) datasets. The predictor-gene signatures (predictor variables; gene*_i_*) and the associated coefficients (coef) were used to construct the molecular prognostic score (mPS) or risk score using the training dataset as shown in the equation below.

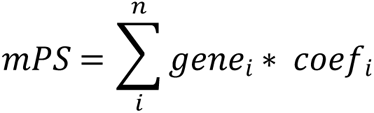

The predictor variables (e.g. genes) and the associated coefficients were further applied to predict the test datasets. Receiver operating characteristic (RoC) curve analyses were performed at a different time-points (in years) for survival data [21] to study predictive capacity.

### The molecular prognostic score (mPS) determines the risk score for over-all survival

The risk scores obtained as mentioned above were used to divide or partition the samples (HGSOC patients) into high (values above median) vs. low-risk groups based on the median values of mPS. The HGSOC samples were also portioned into quartiles or four equal parts based on associated mPS values. Kaplan-Meier survival plot was generated using R with ‘survival’ and ‘survminer’ packages.

### Gene enrichment analysis using GO and KEGG databases

Gene enrichment analysis [22] was done by applying Bioconductor package ‘limma’ [23] to know the role of various pathways associated with different groups in HGSOC cohorts. These functions (goana, kegga) perform over-representation analyses for Gene Ontology (GO) terms or Kyoto Encyclopedia of Genes and Genomes (KEGG) pathways. Here, the list of differential expressed genes (FDR*<*0.05) with the associated Entrez Gene IDs were used as gene set for over-representation or pathway enrichment analysis. The MArrayLM method extracted the gene sets automatically from a linear model fit object [23] The top 20-dysregulated pathways based on p-values were shown.

### Data mining and analyses

The data was retrieved from the data repositories (TCGA, GEO) and analyzed using R/Rstudio: R version 4.1.0 (2021-05-18) and the analysis code and detailed packages and approach can be obtained from the link: https://rpubs.com/siddik/mPS. The various packagesand other associated base packages were described briefly in Appendix A Supplementary file.

## Results

### Differential gene expression between ovarian carcinoma and normal ovarian tissue

A total of 10,225 genes and 1,016 samples with a minimum gene expression matrix correlation of 0.7 were chosen across the five datasets as mentioned in the Methodology section. The detailed information about the samples can be found in Appendix A Suppl Table S1. This includes ovarian cancer samples (n=988) and ovarian surface epithelial cells without any indication of ovarian tumor represented as normal samples (n=28). Multidimensional scaling plots of distances between gene expression profiles of the samples were plotted. The 500 top variable genes among the samples were used to calculate pairwise distances between samples. We observed that samples were either separated or clustered in groups. The samples belonging to the same dataset were clustered together (Appendix A Suppl Fig. S1) indicating the requirement for the removal of batch effect prior to further analysis. The batch effect due to different datasets was removed (Appendix A Suppl Fig. S1) to have a similar pattern of log2-expression ratios among the subjects/ samples irrespective of different datasets. The differential gene expression was performed between normal samples or non-tumor (n=28) vs. primary HGSOC (n=973). Among the analyzed genes, 649 (downregulated) and 473 (upregulated) genes were differentially regulated in the primary HGSOC tumors with respect to ovarian surface epithelial tissues of normal samples (Fig. 2A, and Appendix A Suppl Table S2). The box plots of the top ten dysregulated genes (based on adjusted p-value or FDR) were shown in Fig. 2B. The top ten upregulated genes (with respect to fold change and FDR *<*0.05) were CP (Ceruloplasmin Ferroxidase), FOLR1, TOP2A, CRABP2, MAL, SOX17, CKS2, TPX2, S100A2 and UBE2C. The top down regulated genes in HGSOC were ABCA8, ALDH1A2, BCHE, EFEMP1, NELL2, HBB, TCEAL2, SFRP1, HBA2 and FLRT2. To study the pathways involved in HGSOC, gene enrichment analyses were performed on these 1122 (649+473) differential expressed genes. As per Gene Ontology (GO) database, the upregulated genes (p*<*0.001) were mainly related to cell cycle process, cell cycle transition, cell/nuclear division, chromatin organisation, chromatid segregation, and DNA replication (Appendix A Suppl Table S3). Further, the KEGG pathway analysis showed dysregulation of cell cycle, Complement and coagulation cascades, DNA replication, Oxidative phosphorylation, ECM-receptor interaction, and Drug metabolism - cytochrome P450 (Fig. 2C). The detailed analysis of pathways including the statistics has been shown in Appendix A Suppl Table S4. Since cell cycle related molecules is often found to play important role in cancer, we have further performed the detailed analysis on the molecules involved in this pathway. The Bioconductor package ‘Pathview’: a tool set for pathway based data integration and visualization [24], was used along with downloaded pathway graph data from KEGG pathway database. The differential expressed genes of primary tumor (Appendix A Suppl Table S2) was mapped with the cell cycle related pathway (hsa04110) molecules. Among the 25 molecules that are differentially upregulated in primary HGSOC, 23 molecules are significantly upregulated leading to cell proliferation and tumor mass in HGSOC. The molecules that are found to be significantly upregulated leading to cell cycle events were ARF, Ink4a (CDKN2A), CycD, CycA, CycB, Cdc7, ChK1, MCM (minichromosome maintenance complex component), etc. (Fig. 2D).

**Figure 2:**
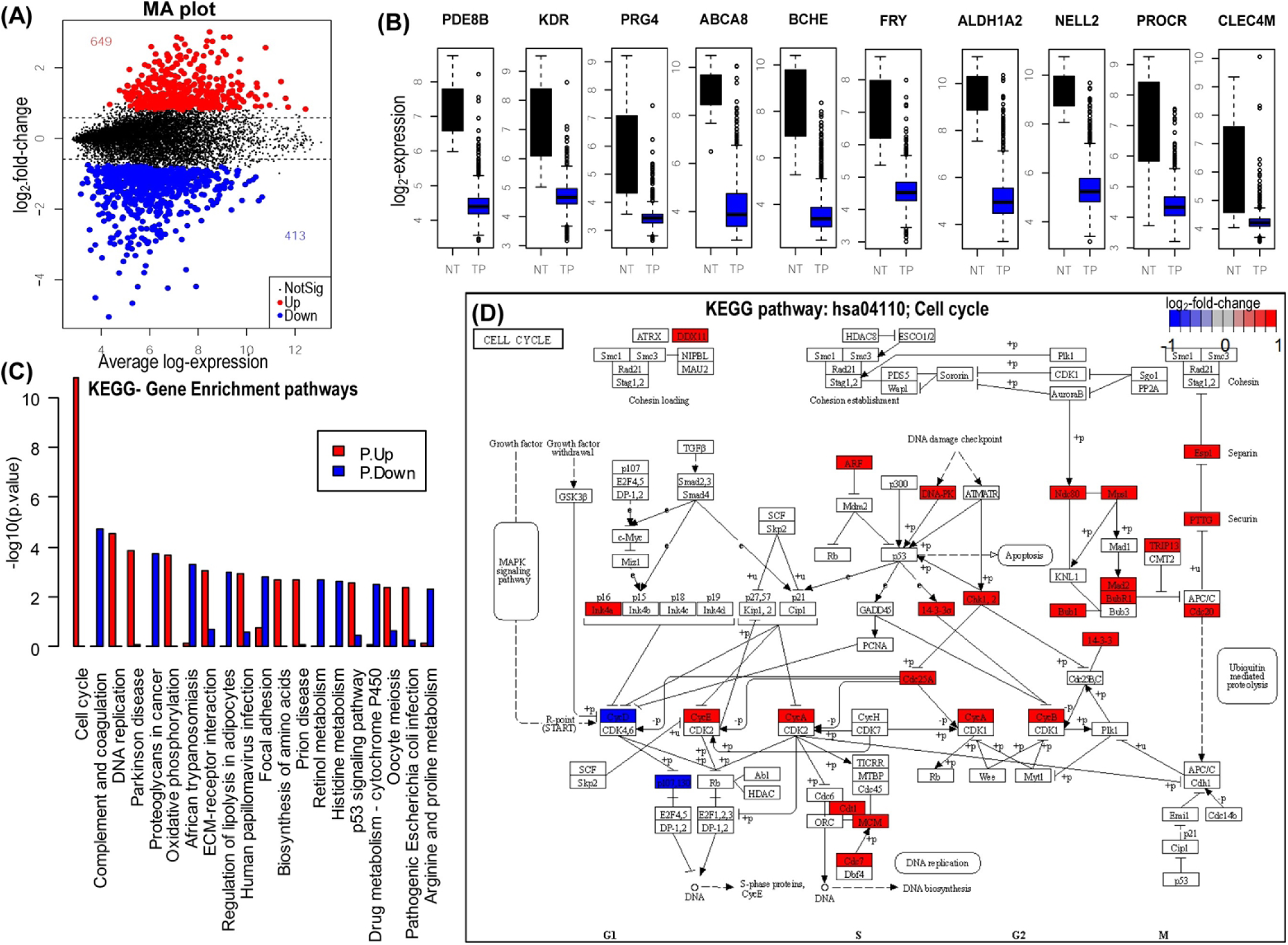
Differential gene expression and pathways involved in HGSOC: *A*; Mean-difference plot (aka MA plot) with color coding for highlighted points (genes) that are differentially expressed in primary tumor (TP) as compared to normal samples (NT). *B*; Box plot showing the top ten dysregulated genes in primary tumor (TP) vs. normal samples (NT). *C*; The key biological/ molecular pathways that are upregulated (P.Up; red color) or downregulated (P.Down; blue) are shown by barplot. The pathways indicated are curated from KEGG pathway database. *D*; The key molecules involve in cell cycle (hsa04110) regulation with FDR*<*0.05 and the indicated log2-fold change are shown by gradient color scale.

### Construction of risk model

The differential expressed genes found in tumor with respect to normal samples with FDR (adj.p-value) *<*0.05 (n=1062) (Appendix A Suppl Table S2) were used as variables to conduct univariate cox regression analyses. The genes were further filtered after applying the logrank test (p-value *<*0.05) and the hazard ratio lies below 0.9 or above 1.1 (0.9*>*HR*>*1.1). There were in total 122 genes of which 63 genes were found to be associated with worse overall survival (HR*>*1.1, p-value*<*0.05) and 59 genes associated with better/ improved overall survival (HR*<*0.9, p-value*<*0.05) of HGSOC patients (Appendix A Suppl Table S5). Finally, these pre-filtered 122 genes were used to construct LASSO estimation using the training dataset (randomly chosen samples; n=491 samples) comprising of both TCGA and GSE cohorts. The log(Λ) vs. partial likelihood deviance plot [25] was shown in Fig. 3 with a different set of alpha (*α*) values. The best fit was observed with *α* =1 (LASSO regression model). The 10-fold cross-validation with α= 1 for deriving LASSO estimation was chosen for further subsequent analysis. The dotted vertical lines indicate the corresponding Λ values (primary x-scale) and gene number (secondary x-scale) with minimal deviance (left). The right vertical line indicates the most regularized model with CV-error within 1 standard deviation of the minimum. There were 20 predictor genes and the associated coefficients were obtained using LASSO regression as shown in Table 1. The detailed analysis of these 20-predictor genes along with the relative expression, fold change (tumor vs. normal), hazard ratio (HR) is shown in Appendix A Suppl Table S6. These 20-predictor genes and the associated coefficients were further used to obtain mPS. This mPS score was used to predict the survival of HGSOC patients.

**Figure 3:**
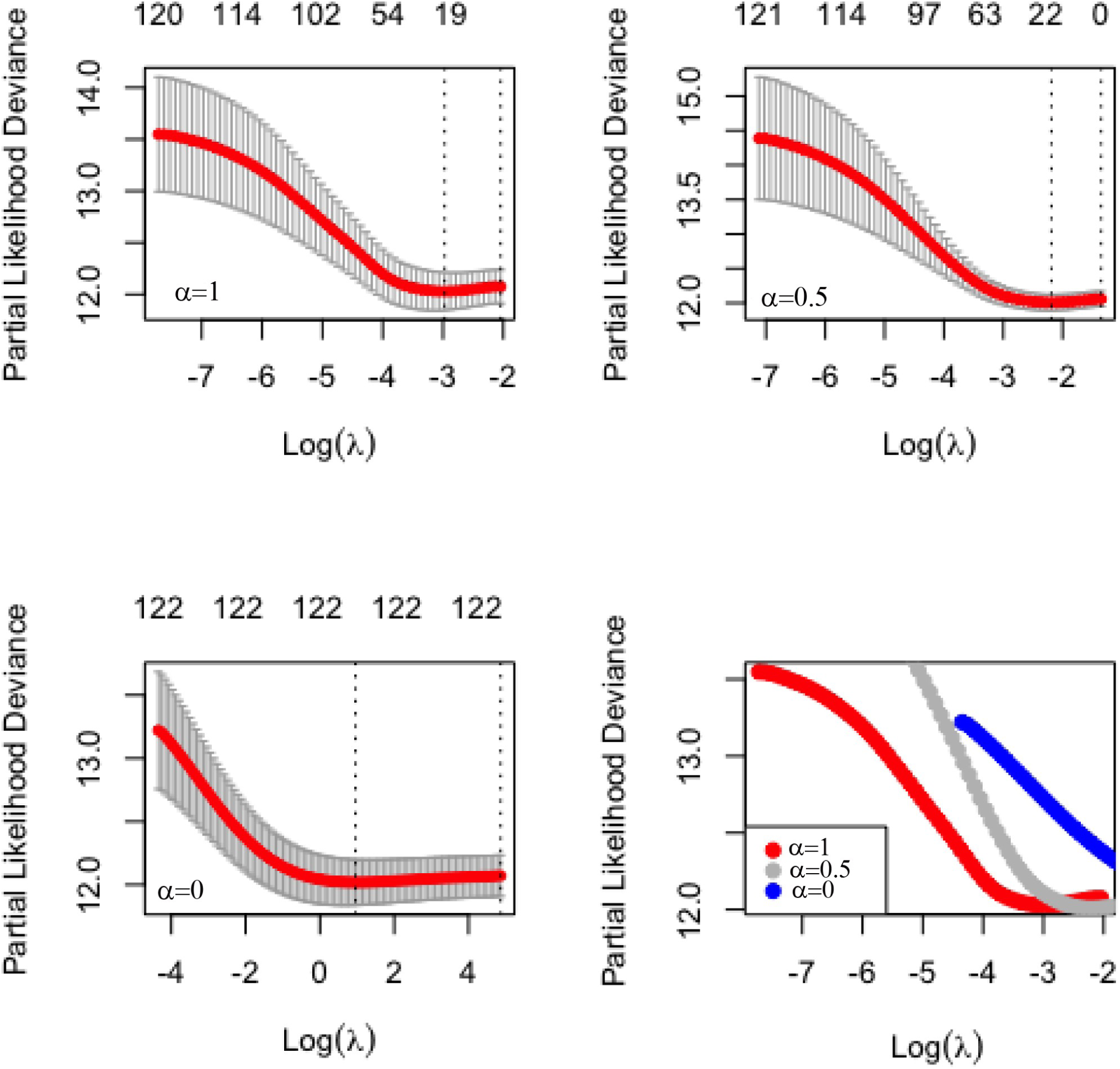
LASSO regression and selection of various parameters: LASSO model fitting on 122 prognostic genes affecting overall survival: The plots with alpha (*α*) = 1, i.e., LASSO (top left), alpha (*α*) = 0.5; elastic net (top right) and alpha (*α*) = 0; ridge regression (bottom left) are shown. The combined/ merged plot (bottom right) with regression curves for LASSO (*α*=1), elastic net (*α*=0.5) and ridge (*α*=0) regression are shown for comparison.

**Table 1:** Predictor genes and associated coefficients.

| SL | gene*i* | coef*i* |
| --- | --- | --- |
| 1 | RHOT1 | 0.1789 |
| 2 | RPS6KA2 | 0.1459 |
| 3 | ASAH1 | 0.1090 |
| 4 | RASA1 | 0.1043 |
| 5 | EDNRA | 0.0750 |
| 6 | NUCB1 | 0.0374 |
| 7 | GFPT2 | 0.0296 |
| 8 | LYVE1 | 0.0201 |
| 9 | PIK3R1 | 0.0149 |
| 10 | BACE2 | -0.0047 |
| 11 | WT1 | -0.0074 |
| 12 | ZNF330 | -0.0075 |
| 13 | GREB1 | -0.0166 |
| 14 | SCTR | -0.0402 |
| 15 | FAM8A1 | -0.0455 |
| 16 | INPP1 | -0.0597 |
| 17 | DIAPH2 | -0.0826 |
| 18 | P2RX7 | -0.0907 |
| 19 | BTN3A3 | -0.1002 |
| 20 | TMED10 | -0.2182 |

### Survival analysis based on molecular prognostic risk score (mPS)

The risk score or molecular prognostic score (mPS) was constructed based on 20 predictor genes. This score was divided into two groups based on median values; high vs low-risk group. The survival or Kaplan-Meier plot was generated as shown in Fig. 4. The training datasets (n=491) and the remaining samples (n=491) were used as test datasets for validation of the mPS by applying the predictor genes and the associated coefficients. The mPS scores at a medium point equally divides the score into higher mPS (higher risk group) and lower mPS (lower risk group). The log-rank p-value (*<*0.0001) of both training and test datasets indicated significant differences in the survival curve of high vs low-risk groups of HGSOC patients. The median overall survival (OS) time (95% lower confidence limit (LCL) - 95% upper confidence limit (UCL)) of high and low-risk groups in the training dataset were 1024 (914-1168) and 1699 (1446-2012) days respectively as shown in Fig. 3A and Table 2. Similarly for the test dataset, the median OS time in days were 1091(1006-1234) and 1976 (1764-2279) for high and low risk groups of HGSOC patients respectively (Fig. 4B and Table 2).

**Figure 4:**
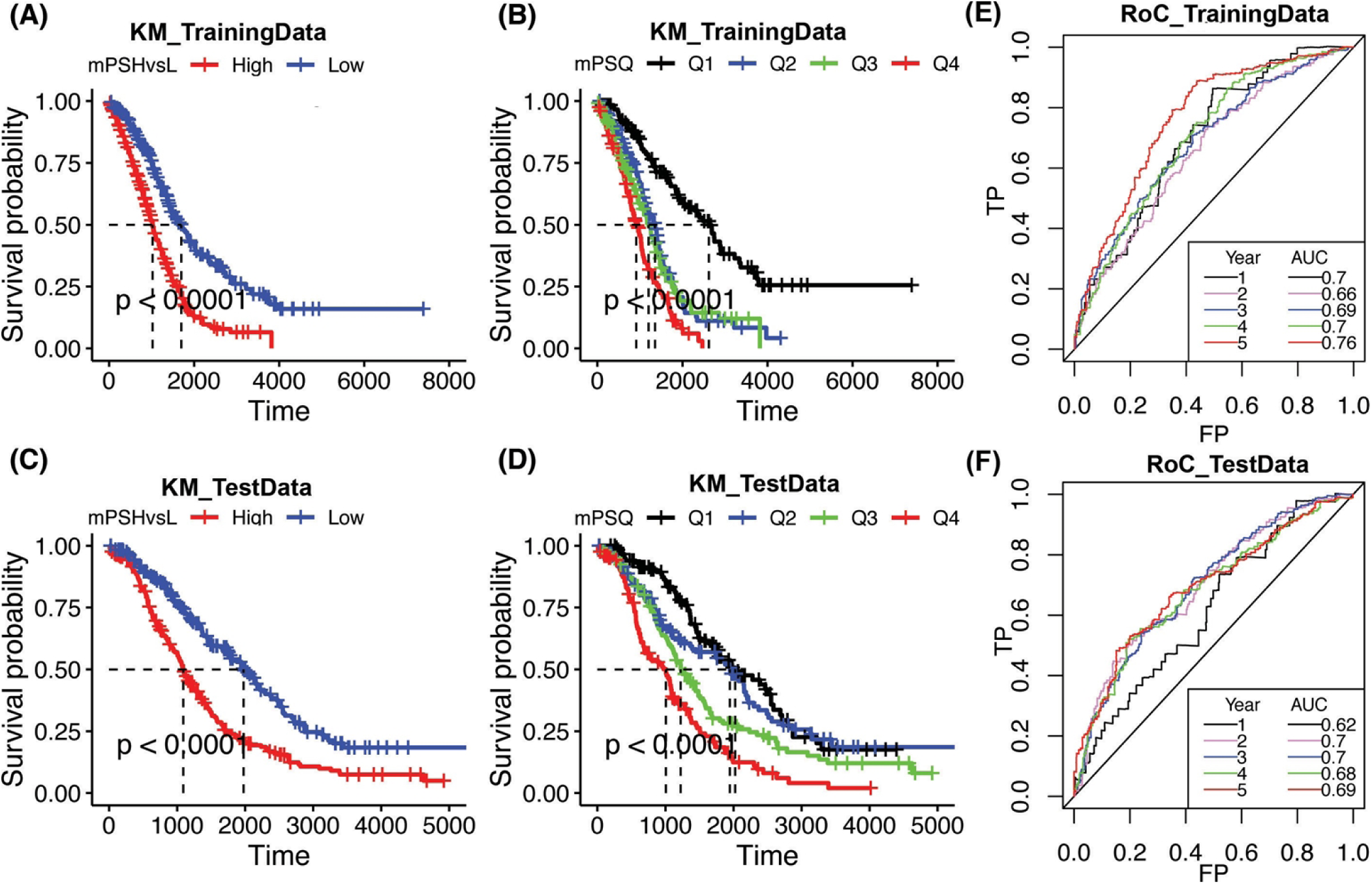
Survival curve and prediction based on mPS score: The cut-off set at median value of mPS (High vs Low mPS) indicates that higher mPS is associated with poor OS time whereas lower mPS is associated with higher OS time (in days) both in training (*A*) and test (*B*) datasets. The groups based on quartiles (Q1; 0-25*^th^* percentiles, Q2; 25-50*^th^* percentiles; Q3; 50-75*^th^* percentiles and Q4; 75-100*^th^* percentiles) showed mPS with higher Q value showed poor OS as opposed to lower Q values both in training (*C*) and test datasets (*D*). The RoC of sensitivity/specificity of test data (*E*) and training data (*F*) for indicated time (in year)is also plotted. TP; True positive, FP; False positive, AUC; Area under curve.

**Table 2:** Groups based on molecular prognostic score and associated median survival.

| Training set |  |  |  |  |  |
| --- | --- | --- | --- | --- | --- |
| Groups | n | events | median | 0.95LC | 0.95UCL |
| High | 245 | 181 | 1024 | 914 | 1168 |
| Low | 245 | 139 | 1699 | 1446 | 2012 |
| Q1 | 123 | 60 | 2621 | 2025 | 3224 |
| Q2 | 123 | 80 | 1354 | 1113 | 1451 |
| Q3 | 122 | 80 | 1203 | 972 | 1389 |
| Q4 | 123 | 101 | 914 | 790 | 1058 |
| Test set |  |  |  |  |  |
| High | 245 | 190 | 1091 | 1006 | 1234 |
| Low | 245 | 131 | 1976 | 1764 | 2279 |
| Q1 | 123 | 62 | 2025 | 1738 | 2553 |
| Q2 | 123 | 70 | 1947 | 1392 | 2218 |
| Q3 | 122 | 91 | 1224 | 1100 | 1484 |
| Q4 | 123 | 99 | 1006 | 687 | 1092 |

Further, we have divided the training samples into four equal parts (quartiles) or subgroups: Q1 (*<* 25*^th^* empirical percentiles), Q2 (25*^th^* to 50*^th^* empirical percentiles), Q3 (50*^th^* to 75*^th^* empirical percentiles), and Q4 (*>* 75*^th^* empirical percentiles) based on the quartiles as cut-off points of mPS score. Q4 has the highest mPS, risk score, followed by Q3, Q2, and Q1. The survival curves of these equally divided quartiles (Q1, Q2, Q3 and Q4) were generated to obtain median over-all survival (OS) time. The median OS time in days were 2621, 1354, 1203, and 914 for Q1, Q2, Q3, and Q4 subgroups, respectively (Fig. 4B, and Table 2). The median mPS of respective quartiles were obtained. Then Pearson correlation between the median mPS of respective quartiles and the respective median OS time was applied. In training data, there was an inverse relationship (r2= -0.902, p=0.049, Pearson correlation) between mPS (risk score) and median OS time which indicates that the mPS score is not only a qualitative indication of survival time but can quantitatively measure or predict the survival time (Fig. 4C and D and Table 2). Similarly, a strong inverse correlation was also obtained for test data (r2=-0.954, p=0.02) between our mPS score and OS time. The heat map as generated using the relative expression of poor predictor (n=9 genes) vs good predictor (n=11) genes (Appendix A Suppl Fig. S2) could potentially cluster both the training and test datasets based on mPS.

### Prediction based on risk score obtained using 20-gene signature

The 20-gene signature was obtained based on values plotted in the graph (Fig. 3) using 10-folds cross-validation of both training and test datasets containing HGSOC samples of different datasets. The derived mPS based on these 20 genes was further applied to study the sensitivity/ specificity using receiver operating characteristic (RoC) curve for survival data. The Area under curve (AUC) values of RoC curves showed the predictive capacity of the prognostic model. The ovarian cancer OS prediction using our prognostic model seemed to be efficient as the AUC values were around 0.70 (±0.03) and 0.68 (±0.03) across the span of 5 years for training HGSOC samples (Fig. 4E) and test data cohorts (Fig. 4F), indicating a very efficient predictor for determining the risk or OS time in HGSOC patients (Fig. 4).

The clinical parameters often used are FIGO stage, tumor grade, residual disease along with age and ethnicity to study the OS time or prognosis of HGSOC. These parameters were converted or scaled into numeric values as shown in Appendix A Suppl Tables S7-S9. Univariate analysis using Cox regression on the survival data of HGSOCs showed that the age, FIGO stage and residual disease at the largest nodule showed a positive correlation (*β* coefficient *>*1; HR *>*1.2, p-value *<*0.05) indicating that the higher values of these parameters showed worse survival or poor prognosis (Table 3). Multivariate Cox regression analysis was performed to adjust for the impact of other significant parameters (Table 3) for deriving the adjusted-HR. Forest plot for themultivariate Cox-proportionalhazards regression model of these parameters was shown in Fig. 5A. The residual disease at the largest nodule showed a significant effect (p*<*0.001) in determining prognosis with an adjusted HR of 1.3 (95% CI=1.13 - 1.40). It inferred that the increasing size of residual disease after cytoreductive surgery was associated with the worse survival of HGSOC patients. Interestingly, the molecular prognostic score (mPS) showed the most significant parameter (p-value *<*0.001) with the HR (adjusted to age, residual disease and FIGO) of 6.1 (95%CI= 3.65 - 10.30) when compared with other clinicopathological parameters along with age.

**Figure 5:**
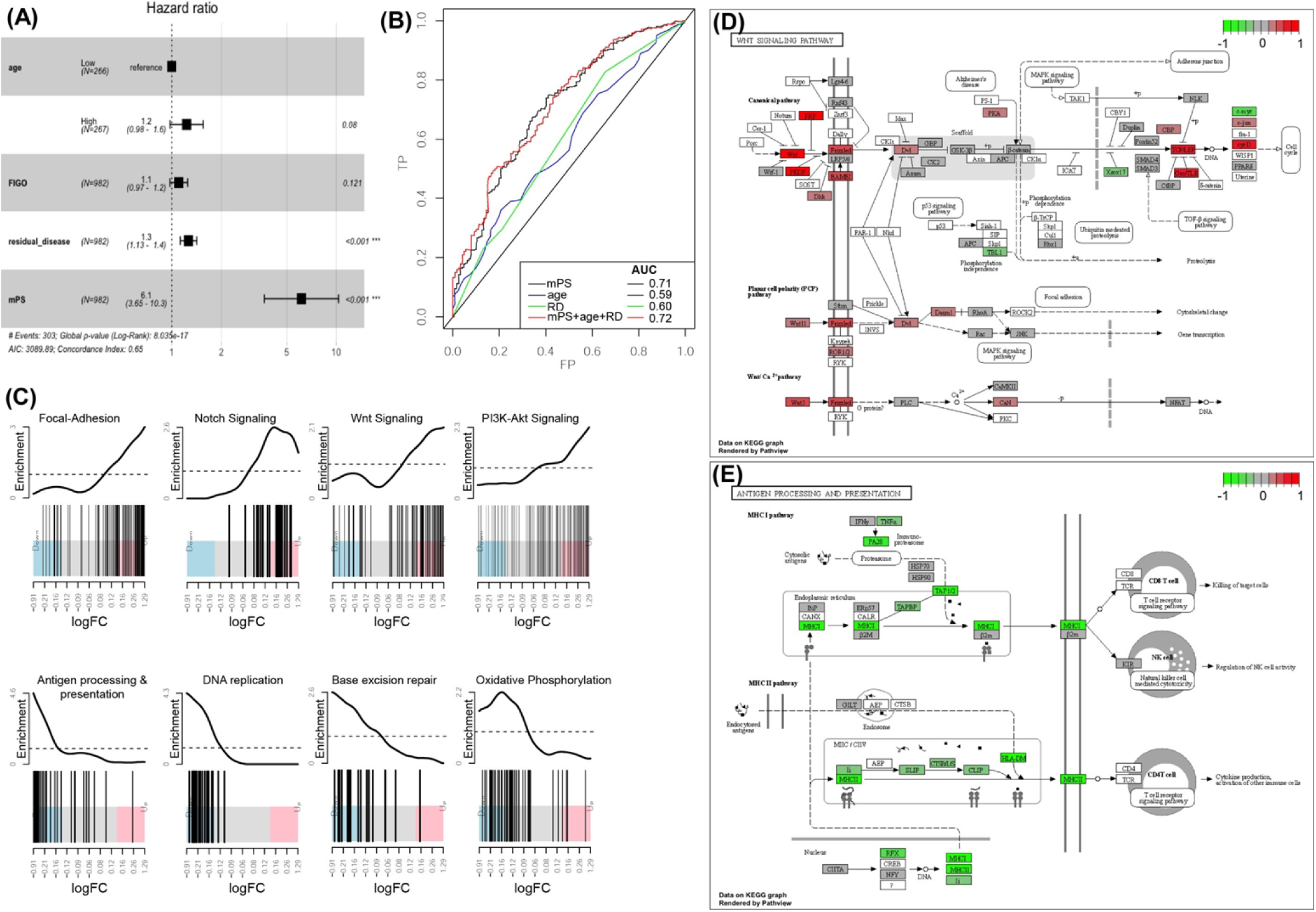
Prognosis of HGSOC using clinicopathological and mPS: *A*; The ggforest plot of Cox proportional Hazard regression fitting of various parameters as indicated. The HR and p-value obtained are adjusted values with respect to other shown parameters. *B;*Area under curve (AUC) using mPS, age and residual disease of the largest nodule (RD) alone or combination as indicated. *C;*Gene enrichment score as shown by barcode plot of indicated KEGG pathways. *D;*Molecules/ genes involved in upregulation of Wnt signaling and *E;* downregulation of Antigen processing and presentation signaling. The log2-fold change of the molecules between high vs low risk group involved are shown with gradient scale.

**Table 3:** Parameters determining prognosis of HGSOC (Univariate Cox regression): The unadjusted-HR, *β* - coefficients are shown here.

| Parameters | $\beta$ | HR (95%CI) | wald.test | p-value |
| --- | --- | --- | --- | --- |
| age at diagnosis (High vs. Low) | 0.32 | 1.4 (1.1-1.7) | 8.5 | 0.0035 |
| FIGO | 0.17 | 1.2 (1.1-1.3) | 8.2 | 0.0041 |
| Tumor grade | 0.21 | 1.2 (0.93-1.6) | 2.1 | 0.15 |
| Ethnicity | -0.1 | 0.9 (0.4-2) | 0.06 | 0.81 |
| Residual disease of largest nodule | 0.28 | 1.3 (1.2-1.5) | 25 | 4.50E-07 |
| molecular Prognostic score (mPS) | 1.9 | 7 (5.1-9.7) | 140 | 3.50E-32 |

We have further analyzed whether the addition of parameters such as age and residual disease of the largest nodule could add prognostic values. It was found that the AUC on the 5-year OS of HGSOC using the mPS score alone was 0.71 as compared to 0.60 contributed by the residual disease of the largest nodule. Moreover, in the mPS score, the addition of parameters such as residual disease and age showed a very nominal improvement in the predictive capacity (AUC= 0.72) of HGSOC patients (Fig. 5B). Thus, mPS score outperformed various traditional parameters, such as age and residual disease of the largest nodule, in terms of prediction of OS of HGSOC. In conclusion, the mPS score alone could serve as a pivotal prognostic factor in predicting the outcome of the severity of HGSOC in terms of OS.

### Gene enrichment study/pathway analysis using high (poor) vs. low (good) risk group

To check the changes in the gene expression between the high risk (having higher mPS) vs. low-risk, differential gene expression was studied. We found that there were 1988 and 2453 significantly (FDR *<*0.05) up and down-regulated genes respectively, in the high-risk group as compared to the low-risk group (Appendix A Suppl Table S10). To check whether there was an involvement of particular pathways or events responsible for the poor survival of HGSOC patients, we performed gene enrichment studies. Gene enrichment by GO related terms indicated the significant (p *<*0.05) downregulation of DNA repair, respiratory electron transport chain, cell cycle, DNA replication related pathways. The cell migration, extracellular matrix interactions, vasculature and blood vessel development were upregulated (p *<*0.05) as shown in Appendix A Suppl Table S11. Similarly, pathway analysis using KEGG pathway database showed similar results. Here we found that Focal-adhesion, Notch signaling, Wnt signaling, PI3-Akt signaling, and signaling pathways regulating pluripotency of stem cells were upregulated (Fig. 5C, Table 4) whereas pathways involving the antigen processing and presentation, cell cycle, DNA replication, and base excision repair were downregulated (Fig. 5C). Since molecules involved in Wnt Signaling [26] as well as the antigen processing and presentation[27] has been reported previously in related to their prognostic importance, we have further investigated or decipher the molecules regulating these two pathways (Fig 5D and E). The Frizzled related family of proteins (FRP) such as FRZB, SFRP1, SFRP4, Wnt family members (WNT4, WNT5A, WNT7A and WNT11), pigment epithelium-derived factor (PEDF), serpin family F member 1 (SERPINF1), Frizzled (FZD) proteins (FZD1, FZD2 and FZD7), BMP and activin membrane-bound inhibitor (BAMBI), segment polarity protein dishevelled (Dvl), protein kinase A (PKA), *β*-catenin, transcription factor-like (TCF)/ lymphoid enhancer-binding factor (LEF), cyclin D1/D2 (cyc-D) were found to be upregulated in the high risk group of HGSOC. This in-lieu leading to the activation canonical Wnt signaling, eventually resulting in cell movement and proliferation (Fig. 5D). Considering the favorable outcome of immunoreactive subtypes [27] in ovarian cancer, we have studied in-detailed, the molecules involved in antigen processing and presentation signaling. This pathway was found to be downregulated (p *<*1.27E-10) in high risk HGSOC patients. There were almost 37 molecules significantly (FDR *<*0.05) downregulated in this pathway. Some of the key mediators such as IFN-*γ*, TNF-*α*, immuno proteasome activator PA28, TAP1/2, TAPBP, MHC-I (HLA-A, HLA-B, HLA-C) were downregulated, affecting MHC-I pathway mediated killing of cancer cells. MHC-II pathway via HLA-DMA, HLA-DMB, HLA-DOA, CLIP (CD74), cathepsin S (CTSS) were also found to be downregulated leading to decreased antitumor cytokine production and activation of other immune cells. Hence, immune evasion and escape was associated with the high or increased risk group of HGSOC patients based on our findings.

**Table 4:** Pathways upregulated in higher mPS (higher risk group) with relative to lower mPS (lower risk group). The details of the pathways are curated from KEGG pathway database. ^1^Total number of molecules involved with the associated KEGG term or pathway; number of differential expressed genes that are ^2^upregulated or ^3^downregulated; ^4^p-value for over-representation of KEGG term in upregulated genes; ^5^p-value for over-representation of KEGG term in downregulated genes.

| Pathway ID | Pathway | N<sup>1</sup> | Up<sup>2</sup> | Down<sup>3</sup> | P.Up<sup>4</sup> | P.Down<sup>5</sup> |
| --- | --- | --- | --- | --- | --- | --- |
| hsa04510 | Focal adhesion | 176 | 79 | 16 | 8.68E-15 | 1.00E+00 |
| hsa04612 | Antigen processing and presentation | 58 | 2 | 37 | 1.00E+00 | 1.27E-10 |
| hsa01100 | Metabolic pathways | 1071 | 137 | 340 | 1.00E+00 | 6.15E-10 |
| hsa04360 | Axon guidance | 149 | 60 | 17 | 2.92E-09 | 1.00E+00 |
| hsa05206 | MicroRNAs in cancer | 145 | 58 | 20 | 7.17E-09 | 9.99E-01 |
| hsa03030 | DNA replication | 32 | 0 | 23 | 1.00E+00 | 1.39E-08 |
| hsa00190 | Oxidative phosphorylation | 88 | 6 | 45 | 1.00E+00 | 2.96E-08 |
| hsa05200 | Pathways in cancer | 455 | 134 | 87 | 1.01E-07 | 9.95E-01 |
| hsa05330 | Allograft rejection | 32 | 1 | 22 | 9.99E-01 | 1.04E-07 |
| hsa01200 | Carbon metabolism | 96 | 4 | 46 | 1.00E+00 | 2.57E-07 |
| hsa05205 | Proteoglycans in cancer | 181 | 64 | 31 | 3.02E-07 | 9.90E-01 |
| hsa04932 | Non-alcoholic fatty liver disease | 135 | 22 | 59 | 8.51E-01 | 3.26E-07 |
| hsa04512 | ECM-receptor interaction | 76 | 34 | 4 | 4.24E-07 | 1.00E+00 |
| hsa05208 | Chemical carcinogenesis - reactive oxygen species | 165 | 29 | 68 | 7.58E-01 | 6.25E-07 |
| hsa05332 | Graft-versus-host disease | 32 | 2 | 21 | 9.91E-01 | 6.82E-07 |
| hsa05415 | Diabetic cardiomyopathy | 154 | 29 | 64 | 6.09E-01 | 9.37E-07 |
| hsa04520 | Adherens junction | 63 | 29 | 7 | 1.45E-06 | 9.97E-01 |
| hsa04010 | MAPK signaling pathway | 259 | 82 | 35 | 1.48E-06 | 1.00E+00 |
| hsa05414 | Dilated cardiomyopathy | 83 | 35 | 5 | 1.60E-06 | 1.00E+00 |
| hsa04151 | PI3K-Akt signaling pathway | 296 | 91 | 40 | 1.65E-06 | 1.00E+00 |
| hsa04310 | Wnt signaling pathway | 133 | 49 | 21 | 1.89E-06 | 9.92E-01 |
| hsa04933 | AGE-RAGE signaling pathway in diabetic complications | 94 | 38 | 18 | 2.03E-06 | 8.92E-01 |
| hsa04810 | Regulation of actin cytoskeleton | 176 | 60 | 31 | 2.80E-06 | 9.84E-01 |
| hsa05012 | Parkinson disease | 198 | 30 | 76 | 9.52E-01 | 3.73E-06 |
| hsa00020 | Citrate cycle (TCA cycle) | 28 | 2 | 18 | 9.82E-01 | 6.76E-06 |
| hsa04926 | Relaxin signaling pathway | 109 | 41 | 17 | 7.09E-06 | 9.88E-01 |
| hsa04919 | Thyroid hormone signaling pathway | 109 | 41 | 16 | 7.09E-06 | 9.94E-01 |
| hsa05169 | Epstein-Barr virus infection | 181 | 23 | 69 | 9.94E-01 | 1.35E-05 |
| hsa05224 | Breast cancer | 127 | 45 | 27 | 1.58E-05 | 7.95E-01 |
| hsa01522 | Endocrine resistance | 84 | 33 | 18 | 1.92E-05 | 7.48E-01 |
| hsa04015 | Rap1 signaling pathway | 178 | 58 | 23 | 1.94E-05 | 1.00E+00 |
| hsa03440 | Homologous recombination | 32 | 0 | 19 | 1.00E+00 | 1.97E-05 |
| hsa04330 | Notch signaling pathway | 45 | 21 | 2 | 3.13E-05 | 1.00E+00 |
| hsa04940 | Type I diabetes mellitus | 38 | 1 | 21 | 1.00E+00 | 3.28E-05 |
| hsa04916 | Melanogenesis | 86 | 33 | 10 | 3.39E-05 | 9.99E-01 |
| hsa01240 | Biosynthesis of cofactors | 99 | 8 | 42 | 1.00E+00 | 3.75E-05 |
| hsa04350 | TGF-beta signaling pathway | 83 | 32 | 19 | 3.97E-05 | 6.35E-01 |
| hsa05165 | Human papillomavirus infection | 285 | 83 | 59 | 4.35E-05 | 9.19E-01 |
| hsa04974 | Protein digestion and absorption | 73 | 29 | 7 | 4.74E-05 | 1.00E+00 |
| hsa03410 | Base excision repair | 31 | 2 | 18 | 9.90E-01 | 4.96E-05 |
| hsa05020 | Prion disease | 211 | 40 | 76 | 5.99E-01 | 5.06E-05 |
| hsa05014 | Amyotrophic lateral sclerosis | 264 | 41 | 91 | 9.59E-01 | 6.59E-05 |
| hsa04550 | Signaling pathways regulating pluripotency of stem cells | 123 | 42 | 29 | 8.04E-05 | 5.78E-01 |
| hsa03430 | Mismatch repair | 22 | 0 | 14 | 1.00E+00 | 8.73E-05 |
| hsa03010 | Ribosome | 100 | 8 | 41 | 1.00E+00 | 1.17E-04 |
| hsa04261 | Adrenergic signaling in cardiomyocytes | 129 | 43 | 11 | 1.25E-04 | 1.00E+00 |
| hsa03040 | Spliceosome | 85 | 9 | 36 | 9.91E-01 | 1.37E-04 |
| hsa05320 | Autoimmune thyroid disease | 41 | 2 | 21 | 9.98E-01 | 1.43E-04 |
| hsa05164 | Influenza A | 143 | 11 | 54 | 1.00E+00 | 1.50E-04 |
| hsa04923 | Regulation of lipolysis in adipocytes | 50 | 21 | 6 | 2.05E-04 | 9.89E-01 |

## Discussion

The 20-gene signature that were used to develop a <u>m</u>olecular <u>p</u>rognostic <u>s</u>core (mPS) could potentially determine the overall survival of HGSOC patients. The AUC (∼ 0.7) of mPS-based stratification both in TCGA and GEO datasets indicates the role of mPS in influencing the overall survival of cancer patients. The mPS determining the overall survival has been previously [11, 12] demonstrated where the mPS score was found inversely correlated with the survival time of patients. In the recent published work [11], it was shown that 101-predictor genes based mPS showed better prediction than age and stage in the advanced HGSOCs. We have also observed the similar findings (Fig. 5A and B). Interestingly, here we have used a lesser number of genes (i.e. 20) than the published work using 101-predictor genes [11]. The mPS scoring system showed much improved power of prediction in HGSOC cohorts of both TCGA and GEO data repositories than the conventional parameters including the age, FIGO score, etc. The similar predictive capacity in both the test and the training datasets of HGSOC patients (Fig. 3 and 4) were observed. Thus, a 20-gene signature derived mPS score could be a better alternative to predict the outcome or survival of HGSOC patients. The differences in predictor genes that we have obtained than from the published report [11] might be attributed due to i) dissimilarities in pre-filtering or screening approaches and ii) total number of genes used during the initial screening process (*∼*10225 by us as compared to 513 genes by Millstein et al.). Our pre-filtering approach was shown in Fig. 1. Initially, the common genes (10225 genes) available across TCGA and GEO data repositories containing HGSOC mRNA expression profiles were selected. It was further filtered to find differential genes in HGSOC tumor vs. normal samples (ovarian surface epithelial cells without any indication of tumor). These genes were finally pre-filtered by Cox proportional hazards regression (univariate) model prior to LASSO-Cox fitting. Thus, the approach of selecting pre-filtered genes prior to application to the LASSO-Cox fitting model was different in our study as opposed to previous reported study by Millstein et. al. [11]. Moreover, the predictor genes involving 101-prognostic genes [11] were derived from a total set of 513 genes, that were also used for the molecular classification of HGSOC [28]. This selection process [11, 28] might left out many important prognostic genes. This might be the reason for the deviation of obtaining prognostic genes in our study as compared to published study[11]. Since gene signature based molecular prognostic score in HGSOC has been previously reported [11], we compared the mPS score obtained using 101-predictor genes [11] and our 20-predictor genes in the integrated HGSOC TCGA and GEO datasets encompassing 982 patients with expression as well as the survival data. Among the 101-predictor genes [11], 85-genes were found common in the integrated TCGA and GEO datasets that we have used for our study. These 85 genes were used to derive the mPS score based on the associated coefficients of these genes as previously reported[11] and the expression data. The AUC of these 85-predictor genes in determining the OS time that ranged from 0.61 - 0.68 for a period of 1 to 5-year (Fig. 6A). Similarly our derived mPS using our 20-predictor genes (Table 1) in the same integrated TCGA and GEO datasets yielded an AUC in the range of 0.67 to 0.72 for a period of 1 to 5-year (Fig. 6B).

**Figure 6:**
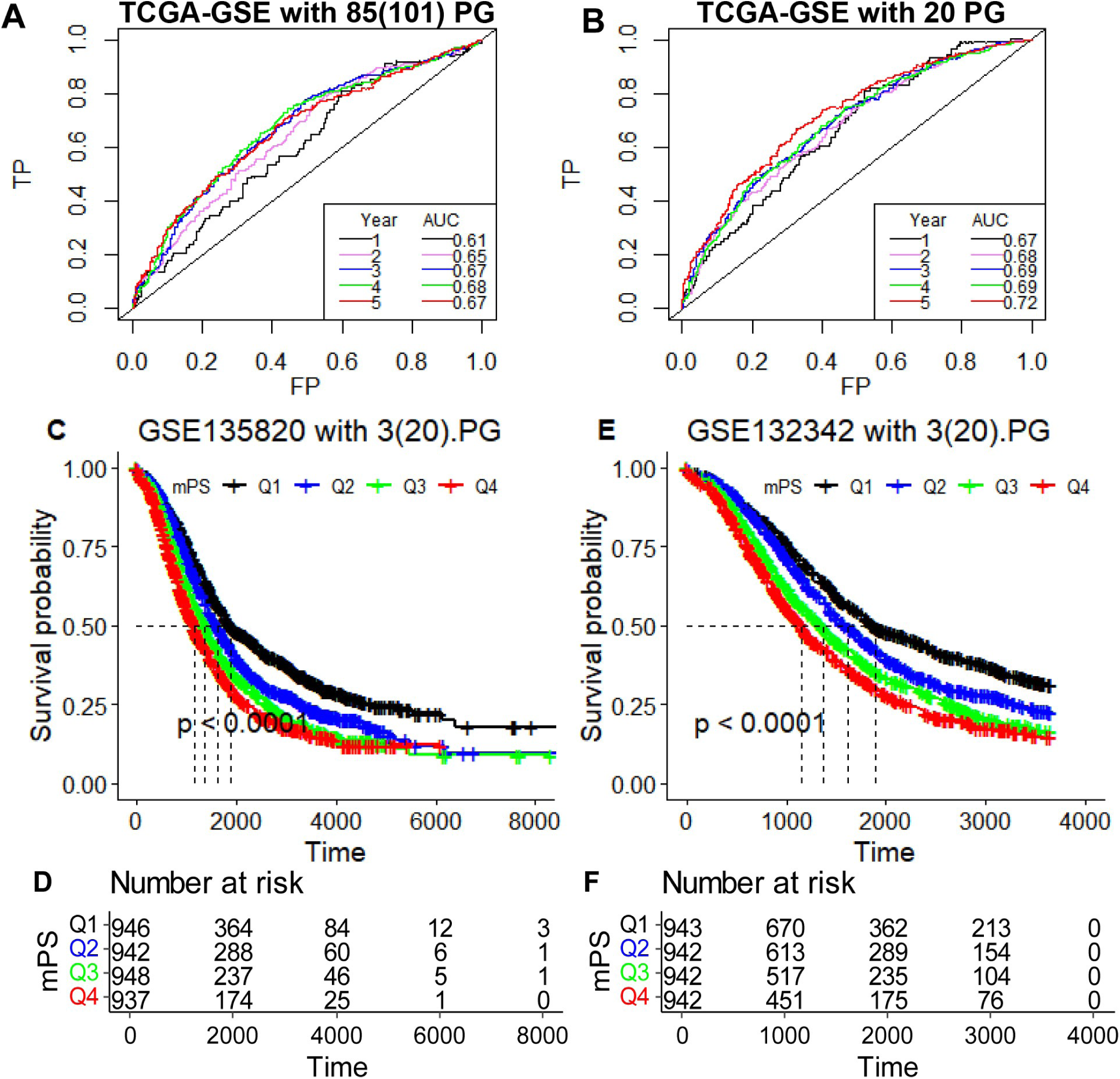
Prognostic performance of molecular prognostic score (mPS)across sequencing platforms and datasets: RoC curves for prognostic performance of mPS derived from the 101-predictor genes *(A)* as described previously [11] along with our (Table 1)20-predictor genes *(B)*. Prediction was studied using AUC for the period of 1 to 5 years’ duration in the integrated datasets (TCGA-OV and GSE14764, GSE18520, GSE26193, GSE26712 and GSE63855) spanning 982 samples. Cross validation across sequencing platforms is done with our 20-predictor genes. There are only 3 out of 20 genes found to be included in the gene expression based NanoString platforms as indicated. Quartiles divides the HGSOC patients into four equal parts based on mPS derived from 3 predictor genes: Q1 bearing the lowest where as Q4 bearing the highest mPS score. Kaplan–Meier curves and the associated risk table of overall survival for HGSOC patients in the GSE135820 (n=3773) *(C, D)* and GSE132342 (n=3769) *(E, F)* datasets.

Similarly, we have applied our 20-gene signature to obtain the mPS score in NanoStringbased RNA sequencing datasets (GSE132342,n=3769;GSE135820, n=3773) that were used in the previous study [11, 28] for across dataset or across platform validation. There were only 3 genes (GFPT2, WT1, RASA1) common between our 20-predictor gene signature and the above mentioned mRNA expression of Ovarian Tumor Tissue Analysis (OTTA) consortium dataset (Appendix ASuppl Table S12). Interestingly, the derived mPS score based on the linear addition of coefficients along with expression of these 3 genes in OTTA dataset (GSE135820, GSE132342) potentially predicted the overall survival of HGSOC patients (Fig. 6 C-F). In-order to study the association the median OS time and mPS, we have partitioned the HGSOC samples into four equal parts based on mPS score; Q1 bearing the lowest mPS value and Q4 bearing the highest mPS value. The median survival time was found to be least with subjects partitioned in the group bearing highest mPS value (Appendix A Suppl Table S13). The median survival time in the groups stratified based on mPS value were found to differ significantly (Fig. 6 C-F).

Our 20-gene expression based molecular prognostic score (mPS) that was derived using Affymetrix human U133A/ U133 Plus 2.0 microarray is a robust dynamic prognostic indicator. The mPS score derived using Affymetrix microarray expression data efficiently predicted the outcome of HGSOC across various datasets. It was also applicable in predicting the outcome of the ovarian cancer irrespective of platforms of mRNA expression data. It efficiently predicted the outcome of ovarian cancer based on the mRNA expression data obtained using NanoString platform (Fig. 6) as well as the Illumina RNASeq platform (Appendix A Suppl Fig. S3).

Apart from the prognostic index of 20-gene signature-based mPS score, a risk classifier, we have in-detailed sttudy the key regulatory pathways that were responsible for the poor prognosis of HGSOCs. It was found that the poor prognostic group or high risk group of HGSOCs have altered pathways regulating TGF-*β* [29], PI3K-Akt [30], Wnt/Notch [31] signaling pathways. These pathways are often associated with poor survival in cancer patients. Immune evasion or escape was also observed in the high risk group, and found to be associated with poor outcome [27]. Thus, targeting these dysregulated pathways [32] might prove beneficial for the high risk group that was anticipated to have poor survival outcomes with current prevailing treatments. Interestingly, we found that the molecules involved in DNA replication and repair, antigen processing and presentation were downregulated in high risk group. Thus, there might be the role of immune evasion or antigenicescape [33] and defective DNA repair pathways [34] in therapy resistance in high risk HGSOC. Hence, further investigation into their role in therapy resistance is needed to find the target molecules and reprogram the HGSOC towards an immune-reactive state. Hence, it’ll be noteworthy to conduct combination therapy using immunotherapies/ agents overcoming immune-suppression and PARP inhibitors [35, 36] in HGSOC patients in an anticipation to improve the overall survival time.

## Conclusion

The conventional parameters like age, clinicopathological parameters: stage/FIGO score, histology, and residual disease showed a trend in prognosis [6], but still the 5-year survival of ovarian cancer remains unchanged. Currently, a molecular gene-based prognostic score derived from the predictor genes and associated coefficients was found to be a robust prognostic marker/ classifier applied in various cancers including breast, prostate and colon cancer. A similar approach was used in ovarian cancer using 101-predictor genes. We have applied only 20-predictor genes to predict the over-all survival of HGSOCs. Our system was found to be universal and robust as it was applicable and reproducible in various gene expression platforms including microarray, RNASeq or NanoString. Our 20-gene signature based-mPS for the prognosis of survival of HGSOC significantly outperformed the conventional parameters: age, residual disease and FIGO score. The high risk group with lower survival time could be benefited by targeted therapies focusing on dysregulated pathways such as TGF-*β*, Notch signaling, DNA repair and antigen processing and presentation pathways.

## Acknowledgements

The authors would like to thank CSIR for providing structural and financial support. Sarbar Ali Saha is the recipient of Council of Scientific and Industrial Research (CSIR)-Junior Research Fellowship (JRF). Siddik Sarkar is the recipient of Start-up Research Grant (SRG/2019/001880) from Science and Engineering Research Board (SERB), INDIA. The results shown here are in part based upon the data (raw file) generated by the TCGA Research Network: https://www.cancer.gov/tcga. The Gene Expression Omnibus (GEO maintained by NCBI): http://www.ncbi.nlm.nih.gov/geo. The authors would like to acknowledge all the contributing laboratories and authors for data repository files associated with following accession number: GSE18520, GSE26712, GSE26193, GSE63885 GSE14764, and GSE135820 along with TCGA-OV cancer project.

## Author contributions

SS designed the study. SAS, PS, SB download the data from data repositories. SS analyzed and interpreted the data. SS, SAS, PS, SB and PM wrote and revised the manuscript. All authors read and approved the final manuscript.

## Data availability/sharing

The relevant data and the supplementary files are shred in the manuscript. The raw data used and/or analysed during the study are available in the TCGA data repository (https://gdac.broadinstitute.org/), and GEO accession number: GSE18520, GSE26712, GSE26193, GSE63885 GSE14764, and GSE135820. The code for R/ Rstudio used for data analysis can be found from the link: https://rpubs.com/siddik/mPS.

## Appendix A. Supplementary data

The following are the Supplementary data to this article and can be found online.

